# A high-performance genetically encoded sensor for cellular imaging of PKC activity *in vivo*

**DOI:** 10.1101/2024.07.19.604387

**Authors:** Takaki Yahiro, Landon Bayless-Edwards, James A. Jones, Lei Ma, Maozhen Qin, Tianyi Mao, Haining Zhong

## Abstract

We report a genetically encoded fluorescence lifetime sensor for protein kinase C (PKC) activity, named CKAR3, based on Förster resonance energy transfer. CKAR3 exhibits a 10-fold increased dynamic range compared to its parental sensors and enables *in vivo* imaging of PKC activity during animal behavior. Our results reveal robust PKC activity in a sparse neuronal subset in the motor cortex during locomotion, in part mediated by muscarinic acetylcholine receptors.

---

Protein kinase C (PKC) is a major mediator of Gq-coupled neuromodulation^1,2^. Although several genetically encoded PKC activity reporters have been developed (Fig. 1a)^3–7^, it remains difficult to image PKC activity *in vivo*. To develop a sensor for *in vivo* studies, we followed the steps from recent efforts for improving other signaling sensors^8,9^. We first compared existing PKC sensors in human embryonic kidney (HEK) 293 cells using two-photon fluorescence lifetime imaging microscopy (2pFLIM). 2pFLIM is advantageous for *in vivo* imaging because it is relatively insensitive to variable expression levels, sample drifting, and wavelength-dependent light scattering^8,10,11^. The maximal responses of these sensors, as elicited by phorbol 12,13- dibutyrate (PDBu) and ionomycin, were relatively small (Fig. 1b–1d and ED Fig. 1). Among these sensors, IDOCKS gave the largest signal, but it involves the overexpression of a functional PKC isozyme. CKAR2 gave the second largest response, and its expression is likely benign because it is a PKC substrate without physiological function. We therefore focused on optimizing the CKAR series sensor.

**Fig. 1.**
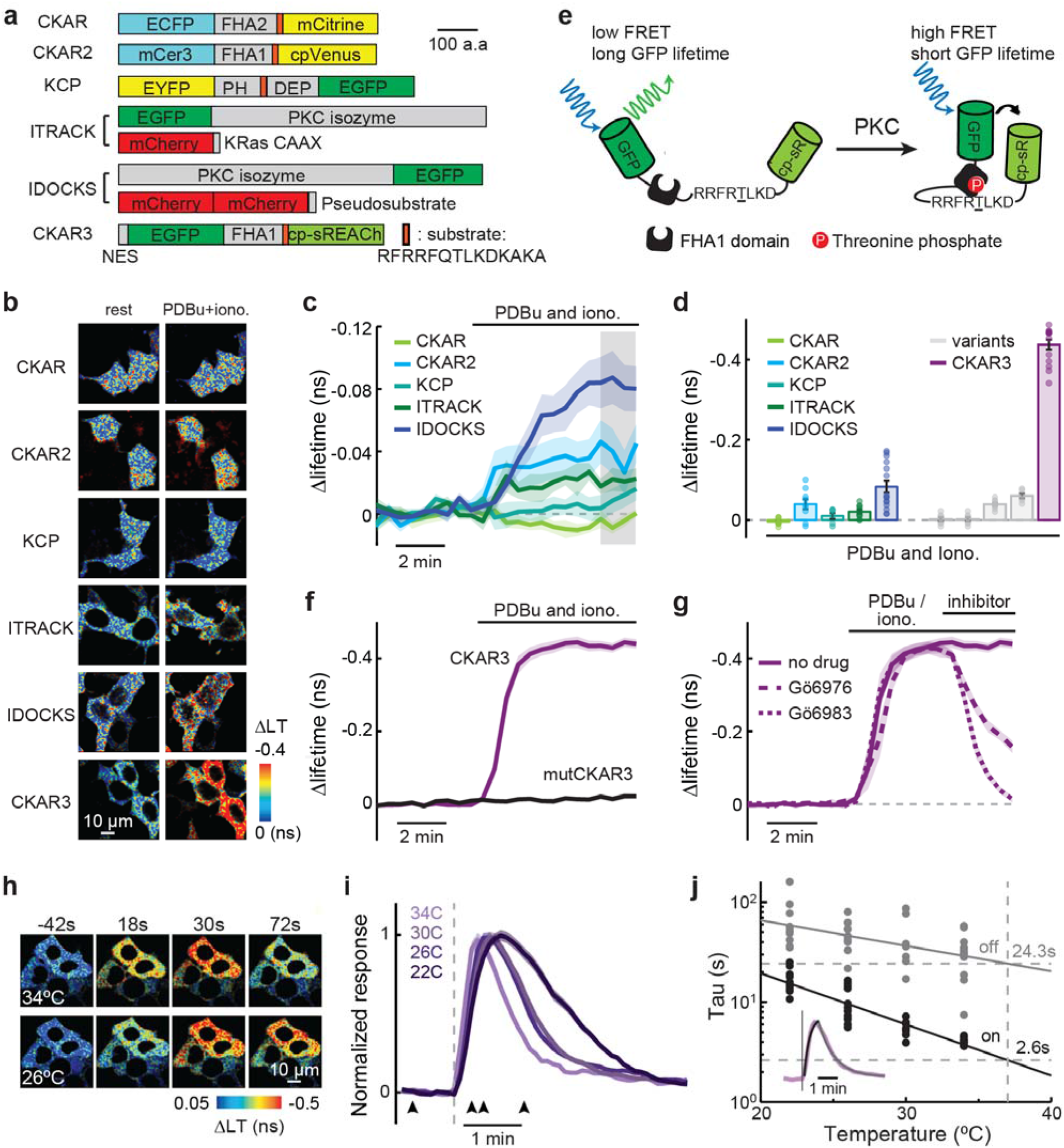
Development and characterization of CKAR3 in cultured cells. **a**, Schematic of previous FRET-based PKC sensors and CKAR3. FHA1 and FHA2: Forkhead-associated domains 1 and 2, respectively. PH: Pleckstrin homology domain. DEP: disheveled, Egl-10, pleckstrin. **b**, Representative Δlifetime images of the tested sensors at rest and after stimulation with 1 μM PDBu and 1 μM ionomycin (iono.). **c**, Averaged response (ΔLT) traces of existing sensors. **d**, Quantification of existing sensors (at the grey window of panel **d**) and candidate variants. From left to right, n (cells/coverslip) = 11/4 9/4, 6/3, 10/5, 15/6, 7/3, 7/3, 10/3, 6/3, and 14/4. **e**, Schematic of CKAR3 sensor. **f**, Response traces of CKAR3 and mutCKAR3 to PDBu and ionomycin. n (cells/coverslips) = 14/4 and 11/4 for CKAR3 and mutCKAR3, respectively. **g**, Responses of CKAR3 to PDBu and ionomycin followed by indicated antagonists. n (cells/coverslips) = 14/4, 7/3, and 8/3 for no drug, Gö6976, and Gö6983, respectively. **h–j**, Representative images (**h**) and traces (**i**) and collective on and off tau (**j**) of CKAR3 responding to hM3D activation at the indicated temperatures. From hot to cold, n (cells/coverslips) = 12/4, 7/4,12/4,11/3. Inset in panel **j**: An example trace and its fittings. Where applicable throughout the figure, mean (dark lines and bars) and s.e.m. (shaded area and error bars) are shown.

CKAR and CKAR2 use the FHA2 and FHA1 phosphopeptide-binding domains, respectively^4,5,12^. Upon phosphorylation of a substrate peptide specific to PKC within the sensor^4,13,14^, the FHA domain binds to the phosphorylated peptide. This changes the sensor protein conformation, thereby altering Förster resonance energy transfer (FRET) between two fluorescent proteins (FPs) at two ends of the sensor. Both CKARs use cyan and yellow FP pairs (Fig. 1a). To increase the sensor dynamic range, we attempted several strategies, including i) using mTurquoise2^15^ as the donor FP to increase the Förster distance (R_0_) of FRET (variants 1– 4) (ED Fig. 2), ii) using tandem-dimer cpVenus to increase FRET efficiency (variants 1 and 3)^16^, and iii) switching the substrate peptide in the lifetime-only PKA sensor, tAKARβ^8^, to that for PKC. The first two strategies resulted in little improvements under lifetime imaging (Fig. 1d).

However, the third strategy greatly enhanced the lifetime signal (Δlifetime = -0.43 ± 0.003; ∼11 fold larger than CKAR2, and ∼5 fold larger than IDOCKS; Fig. 1d). This signal amplitude is comparable to that of the PKA sensor tAKARα, which is sufficient for *in vivo* PKA activity imaging^8^. We call this new sensor CKAR3 (Fig. 1e).

We extensively characterized CKAR3, first in HEK cells. A T-to-A mutation at the phosphorylation site (T416A; construct called mutCKAR3) eradicated the response to PDBu- ionomycin (Fig. 1f). In addition, CKAR3 responses were reverted by bath application of the broad-spectrum PKC inhibitor Gö6983 (1 μM) (Fig. 1g). Interestingly, the PKCα and PKCβ inhibitor Gö6976 only partially reverted the response (Fig. 1g). CKAR3 was also partially activated by PDBu when pretreated with Gö6976 but not with Gö6983 (ED Fig. 3). Together, these results indicate that CKAR3 response is specific to PKC phosphorylation and mediated by multiple PKC isoforms. To determine the sensor kinetics, we co-expressed the Gq-coupled hM3D receptor together with CKAR3. Using short puffs (1-s long) of the ligand C21, we found that the sensor exhibited temperature-dependent on- and off-kinetics with Q10 values of 3.3 and 1.8, respectively (Fig. 1h–1j). By extrapolation, the sensor was estimated to exhibit a τ_on_ of 2.6 ± 1.7 s (fit ± 95% confidence level) and a τ_off_ of 24.3 ± 2.7 s at 37 °C. These values were likely overestimations because it was impossible to use instantaneous puffs, and the stimulant took time to dissipate. Finally, we found that 36% of CKAR3-expressing cells responded to ATP application (25 μM), and these responses were abolished by the Gq-coupled P_2_Y_11_ receptor^17^ antagonist NF-157 (1 μM) (ED Fig. 4), consistent with recent reports^18,19^. Together, these results indicate that CKAR3 responds to Gq activation of PKC.

To test CKAR3’s performance in neurons, we expressed it in CA1 neurons of cultured hippocampal slices using the biolistic transfection methods. CKAR3 exhibited robust responses to acetylcholine (20 μM) in CA1 neurons (Fig. 2a–2c), which is consistent with the finding that these neurons express Gq-coupled muscarinic acetylcholine receptors (mAchRs)^20^. Additionally, CKAR3 responses were much larger than those from CKAR and CKAR2 under the same conditions (Fig. 2a–2c) and were abolished by pretreatment of the broad spectrum mAchR antagonist scopolamine (1 μM) or the T416A mutation (Fig. 2d–2f). Thus, CKAR3 is sufficient for detecting PKC activity driven by mAchRs in CA1 neurons.

**Fig. 2.**
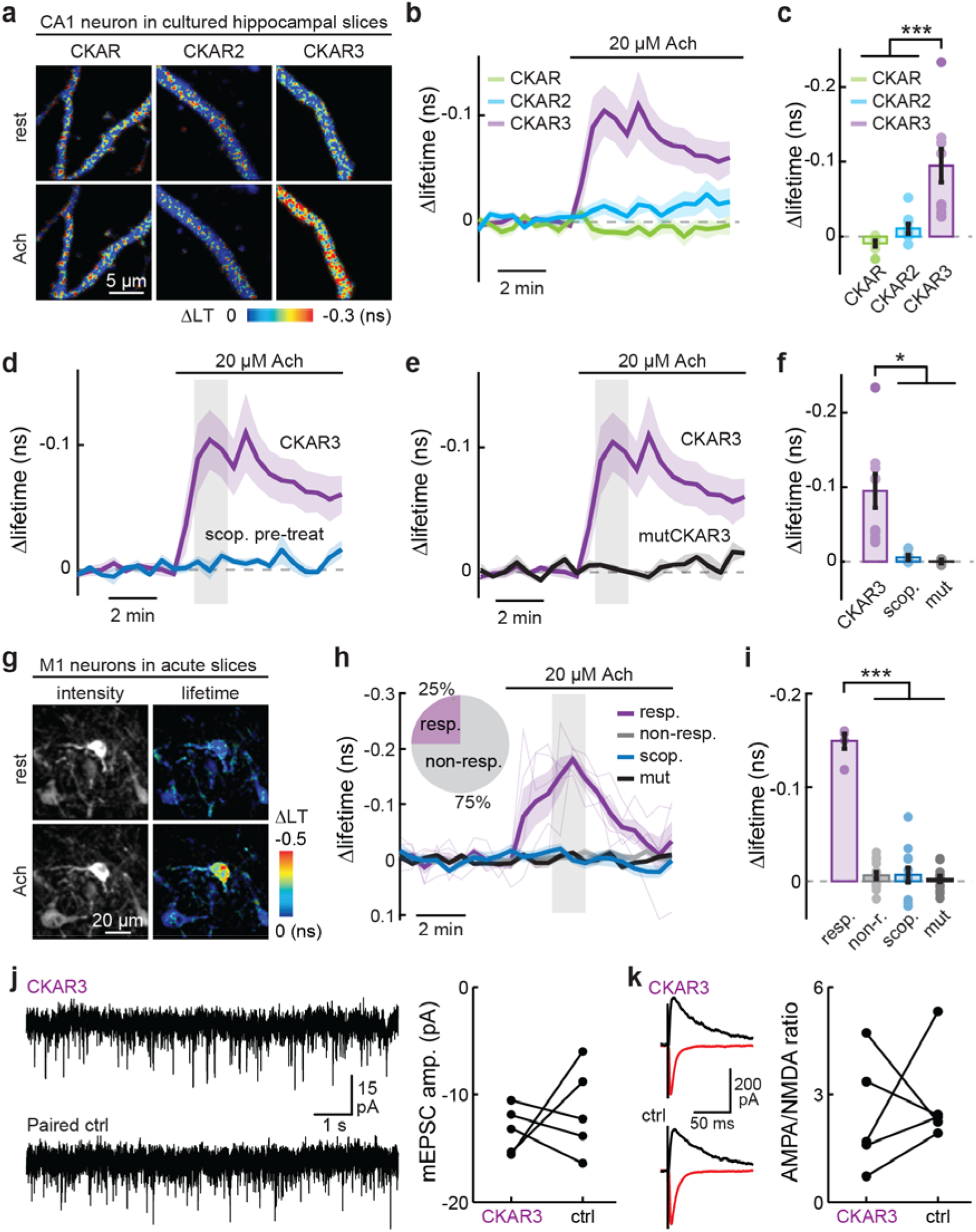
Characterization of CKAR3 in neurons from brain slices. **a**–**c**, Representative images (**a**), average traces (**b**), and quantification (**c**) of CKAR sensors responding to acetylcholine (Ach, 20 μM) in CA1 neurons of cultured slices. n (cells) = 7, 9 and 9 for CKAR, CKAR2 and CKAR3, respectively. ***: p < 0.001 (2x10^-4^ and 1x10^-3^, respectively), dF = 2, one-way ANOVA with Bonferroni post-hoc test. **d**–**f**, Response traces of CKAR3 in CA1 neurons to acetylcholine compared to pre-treatment with scopolamine (scop., 1 μM) (**d**), mutCKAR3 (**e**) and their collective quantifications (**f**). In panel **f** from left to right, n (cells) = 9, 5, and 3. *: p = 0.02 for scopolamine and 0.04 for mutant comparison, dF = 2, one-way ANOVA. **g**, Representative images of CKAR3 in M1 neurons in acute slice in response to acetylcholine. **h** & **i**, Average traces (**h**) and collective quantifications (**i**) of CKAR3 responders (purple; threshold 2x baseline s.d.), non-responders (grey), those with scopolamine pretreatments (blue), or mutCKAR3 responding to acetylcholine application in acute slice. In panel **i** from left to right, n (cells) = 5, 15, 13 and 15. ***: p = 0 for all comparisons, dF = 3, one-way ANOVA. **j**, Representative mEPSC recordings (left) and quantified mEPSC amplitudes (right) from paired adjacent L5 M1 pyramidal neurons with and without CKAR3 expression. n (pairs/mice) = 5/4, p = 0.52, dF = 4, paired t-test. **k**, Representative recordings (left) at holding potential of −70 (red) and +55 (black) mV and quantified AMPA/NMDA receptor current ratios (right) from paired adjacent L5 M1 pyramidal neurons with and without CKAR3 expression. n (pairs/mice) = 5/2, p = 0.79, signed-rank test. Where applicable throughout the figure, mean (dark lines and bars) and s.e.m. (shaded area and error bars) are shown.

To test CKAR3’s responses in cortical neurons, we expressed it in the mouse motor cortex via adeno-associated virus (AAV) injection. Layer 5 (L5) neurons were imaged in acute slices 20–25 days post infection. The sensor expressed well (ED Fig. 5), and it responded to bath-applied acetylcholine in a subset (*∼*25%) of imaged L5 neurons (Fig. 2g and 2f). These responses were abolished by scopolamine or the T416A mutation. In addition, CKAR3 expression did not perturb a palette of tested neuronal properties in these neurons, including the amplitude and frequency of miniature excitatory postsynaptic currents (mEPSCs), the AMPA and NMDA receptor current ratio, and the paired-pulse ratio (PPR) (Fig. 2j and 2k, and ED Fig. 6a and 6b). The intrinsic excitability of cortical neurons was also unaffected (ED Fig. 6c–6j).

We asked whether CKAR3 was sufficient for *in vivo* imaging. The sensor expressed well *in vivo* via AAV injection and could be imaged longitudinally in awake mice in all tested brain regions, including the primary motor cortex (M1), the primary visual cortex (V1), and the lobule VI of the cerebellum (Fig. 3a). Different brain regions exhibited different basal lifetimes, but they were all significantly lower than those of mutCKAR3 (Fig. 3b), corresponding to higher PKC activities. The basal lifetimes of individual neurons were highly correlated week to week (Fig. 3c), suggesting that different brain regions and neurons exhibit distinct set points of basal PKC activity. Animal locomotion elicited robust and repeatable PKC activity responses in a subset (∼31%) of neurons in the M1 cortex but not in the V1 cortex or lobule VI (Fig. 3d–3i) or when mutCKAR3 was used (Fig. 3g and 3h). This sparsity of responders is notably different from that for PKA activities under locomotion, in which a majority of M1 cells responded^8^. The responders could be separated into two distinct clusters (Fig. 3j and ED Fig. 7). The first cluster exhibited a fast-onset, transient response upon the start of locomotion (Fig. 3k). In contrast, neurons in the second cluster were gradually recruited with a delay from locomotion onset, and often exhibited multiple response peaks until the end of locomotion (Fig. 3k and 3l). These results reveal previously unappreciated, cell-specific heterogeneity in PKC signaling. By isolating and averaging the on- and off-responses, we found that both clusters exhibited comparable on- and off-kinetics, with a τ_on_ of ∼4 s and a τ_off_ of ∼14 s (Fig. 3m).

**Fig. 3.**
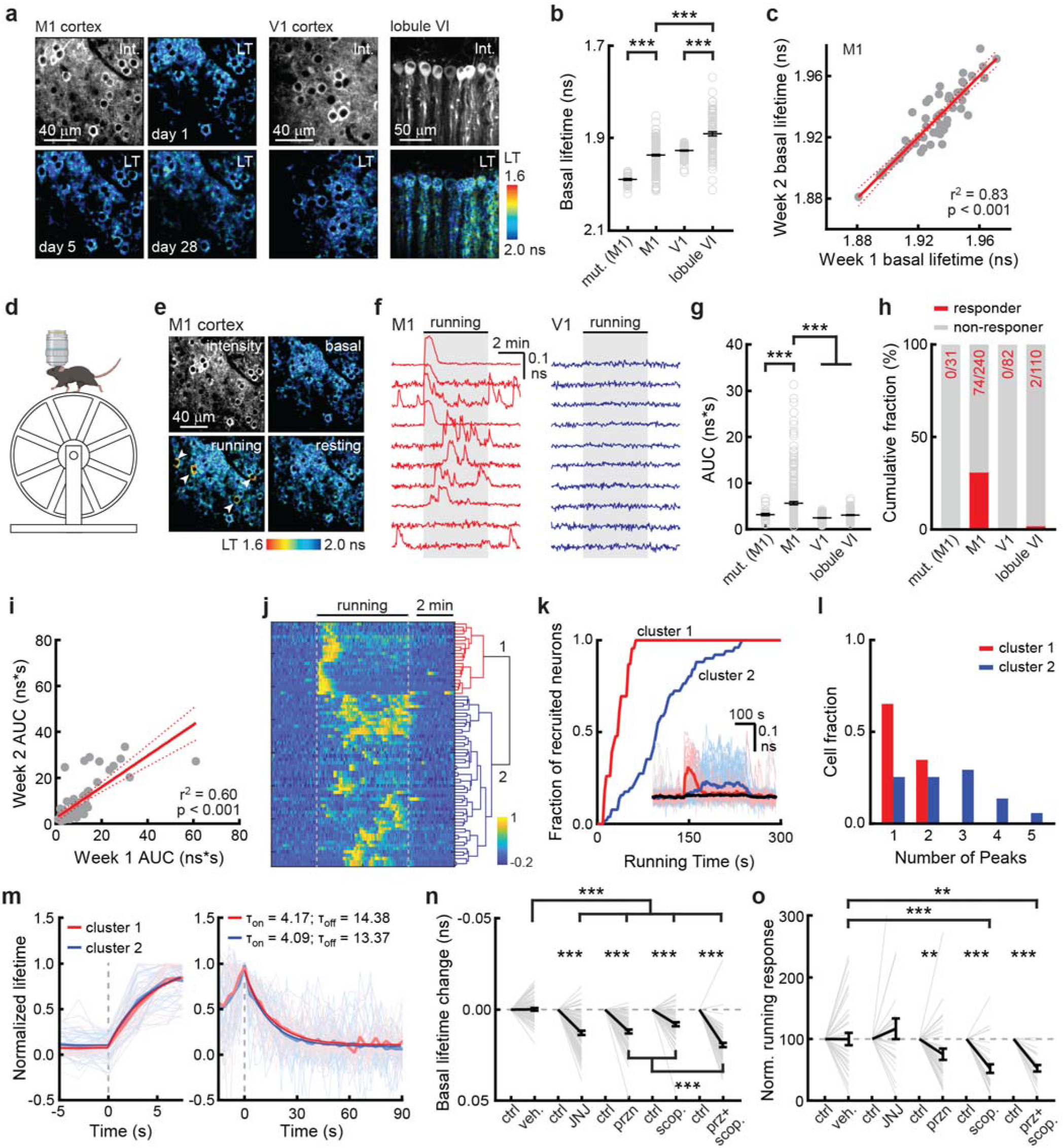
*In vivo* CKAR3 imaging revealed cell-specific PKC activity elicited by behavior. **a** & **b**, Representative intensity and lifetime images (**a**) and quantifications of basal lifetimes (**b**) of CKAR3 or mutCKAR3 in L2/3 neurons of the M1 and V1 cortices and Purkinje cells of cerebellar lobule VI in awake mice, as indicated. LT: lifetime; int.: intensity. In panel **b** from left to right, n (cells/mice) = 31/2, 240/5, 82/3, and 110/3. **c**, Correlation of basal lifetimes with fit and 95% confidence intervals of the same M1 neurons across approximately a week. n (cells/mice) = 65/4. **d**, Schematic imaging during enforced running created in Biorender.com. **e** & **f**, Representative images (**e**) and example traces (**f**) of CKAR3 in L2/3 M1 or V1 neurons during enforced running. **g** & **h**, Quantification (**g**) and fraction of responding cells (**h**, threshold 5x baseline s.d.) of CKAR3 and mutCKAR3 in L2/3 or Purkinje neurons in the indicated brain regions responding to enforced running. Kruskal-Wallis test was used. **i**, Correlation of AUC with fit and 95% confidence intervals in the same L2/3 M1 neurons during enforced running across approximately one week. n (cells/mice) = 74/4. **j**, Hierarchical clustering of the responses of 74 responder neurons. **k**, Cumulative responder fractions over time of each cluster. Inset: average response trace of each cluster. n (cells/mice) = 22/5 for cluster 1, 52 for cluster 2. **l**, Fraction of neurons of each cluster with indicated number of responding peaks during the bouts. **m**, On- and off-kinetics of CKAR3 in L2/3 M1 neurons. *τ*_on_ and *τ*_off_: time constant of on- and off-kinetics. **n** & **o**, Comparison of basal lifetime change (**n**) and response change to enforced running (**o**) induced by indicated drugs. From left to right, n (cells) = 85, 84, 76, 70, 71, 35, 26, 35, 30, and 25 from 4 mice. Note that only responders are included in panel **o**. Wilcoxon test for paired test; Kruskal-Wallis test for comparison across groups. Throughout the figure, horizontal black lines indicate mean and s.e.m.; *: p < 0.05, **: p < 0.01, ***: p < 0.001.

Finally, we interrogated the upstream receptor mechanism of *in vivo* PKC activity using pharmacological manipulations. Basal PKC activity was reduced by systemic injection of antagonists of several Gq-coupled receptors, including metabotropic glutamate receptors (mGluR; JNJ16259685, 4 mg/kg), α1-adrenergic receptors (prazosin, 1 mg/kg), and mAchRs (scopolamine, 1 mg/kg) (Fig. 3n). The effects of antagonizing α1-adrenergic receptors and mAchRs on basal PKC activity were additive, suggesting that the basal activity is controlled by the combined action of multiple neuromodulators. In contrast, locomotion-driven PKC responses were dramatically reduced by the mAchR antagonist scopolamine, but less affected by mGluR and α1-adrenergic receptor antagonists, indicating that different mechanisms regulate basal versus locomotive PKC activities.

Overall, we present a novel PKC activity sensor, CKAR3, which exhibits a >10-fold enhanced dynamic range. 2pFLIM imaging of CKAR3 *in vivo* revealed sparse, but robust, PKC dynamics during animal locomotion that were brain region-specific and largely dependent on mAchR signaling. This sensor should be applicable to PKC activity imaging in many *in vitro* and *in vivo* contexts.

## Methods

All surgical and experimental procedures were performed in accordance with the recommendations in the Guide for the Care and Use of Laboratory Animals, written by the National Research Council (US) Institute for Laboratory Animal Research, and were approved by the Institutional Animal Care and Use Committee (IACUC) of the Oregon Health and Science University (#IP00002274).

### Plasmid Constructs and virus production

Constructs were made using standard mutagenesis and subcloning methods, or by gene synthesis (Genewiz). All previously unpublished constructs and their sequences will be deposited to Addgene. AAV2/1 viruses were packaged by NeuroTools (https://neurotools.virus.works). They express the sensor or mutant either under the human synapsin promotor or, when the Cre-dependent FLEX control is implemented, under a CAG promotor.

### Cell culture and transfection

HEK-293 cells (ATCC #CRL-1573) were maintained in 100-mm cell culture dishes (Fisher Scientific, #FB012924) at 37°C with 5% CO_2_ in minimal essential medium (ThermoFisher #11095-080) with the addition of 10% fetal bovine serum. All cells from ATCC have been authenticated by morphology, karyotyping, and PCR based approaches. These include an assay to detect species-specific variants of the cytochrome C oxidase I gene (COI analysis) to rule out inter-species contamination, and short tandem repeat (STR) profiling to distinguish between individual human cell lines to rule out intra-species contamination. These cells are also tested for mycoplasma by ATCC. Cell aliquots were kept frozen in liquid nitrogen until use, and were further authenticated based on their morphology. Once thawed, each aliquot of cells was passed and used for no more than 4 months. To image transfected HEK cells under our two-photon microscope, cells were subcultured on glass coverslips (hand-cut to ∼5 x 5 mm) coated with 0.1 mg/mL Poly-D-Lysine (Millipore-Sigma, #27964-99-4) in 35-mm cell culture dishes (Corning, #CLS430165). For sensor expression, constructs in mammalian expression plasmids (0.5 µg/35-mm dish unless otherwise noted) were transiently transfected using Lipofectamine-2000 (ThermoFisher, #11668030) according to the manufacturer’s instructions with the exception that only 3 μl of the reagent was used. ITRACK and IDOCKS were transfected with 0.5 µg of the donor construct with at 1:3 donor-acceptor ratio for ITRACK and 1:2 ratio for IDOCKS. For kinetics experiments, cells were transfected with 0.5 µg CKAR3 and 0.25 µg hM3D (i.e., Gq-coupled DREADD). A single-fluorophore PKC sensor ExRai-CKAR^3^, which did not express well in our expression system and was not expected to exhibit lifetime changes, was not included in the comparison. Imaging was performed at two days post-transfection in a chamber perfused with carbogen (95% O_2_/5% CO_2_) gassed artificial cerebral spinal fluid (aCSF) containing (in mM) 127 NaCl, 25 NaHCO_3_, 25 D-glucose, 2.5 KCl, 1.25 NaH_2_PO_4_, 2 CaCl_2_, and 2 MgCl_2_. Drugs were bath-applied with the exception that C21 (0.5 mg/mL) was puffed using a Picospritzer II apparatus (16 psi) for the kinetics experiments.

### Hippocampal slice culture and transfection

Hippocampi were dissected from P6–7 rat pups of both sexes. Sections (400 µm) were prepared using a chopper in dissection medium containing (in mM) 1 CaCl_2_, 5 MgCl_2_, 10 glucose, 4 KCl, 26 NaHCO_3_, and 248 sucrose, with the addition of 0.00025% phenol red. The slices were then seeded onto a cell culture insert (Millipore, #PICM0RG50) and cultured at 35°C with 5% CO_2_ in 7.4 g/L MEM (ThermoFisher, #11700-077) with the addition of (in mM unless labeled otherwise): 16.2 NaCl, 2.5 L-Glutamax, 0.58 CaCl_2_, 2 MgSO_4_, 12.9 D-glucose, 5.2 NaHCO_3_, 30 HEPES, 0.075% ascorbic acid, 1 mg/mL insulin, and 20% heat-inactivated horse serum. Slice media were refreshed every 2-3 days after seeding replacing ∼60% of the culture media.

Transfection of cultured slices was accomplished using the biolistic method (Bio-Rad Helios gene gun) 10 to 20 days after seeding. In short, slices were bombarded with biolistic particles created by coating 1.6 μm gold particles (Bio-Rad, #165-2262; ∼1 μg DNA/mg gold) with constructs in mammalian expression plasmids. Cultured slices were imaged at 2-3 days post-transfection in a chamber perfused with carbogen-gassed aCSF that contained 4 mM CaCl_2_ and 4 mM MgCl_2_.

### Animal surgeries

C57BL/6 mice (both sexes from Charles River or home-bred within 5 generations from Charles River breeders) and *Pcp2-Cre* mice (Jax#010536; both sexes; for cerebellum labeling) were used. Mice were transfected by stereotaxic viral injection, as previously described^21,22^. Cranial window surgeries were performed as previously described^9,23^. Briefly, a circular craniotomy (∼3.5 mm in diameter) was performed using an air-powered high-speed dental handpiece (Maxima Pro 2) and a ¼ inch carbide burr (Henry Schein #5701072) above the region of interest. 100 nL of AAV was then injected into multiple locations within the craniotomy with the following coordinates (in mm anterior to the bregma, lateral to the midline, below the pia): for the M1 motor cortex, (0.0, 1.0, 0.4), (0.5, 1.5, 0.4), and (1.0, 2.0, 0.4); for the V1 visual cortex: (−3.0, 2.5, 0.4), (−3.5, 2.5, 0.4), and (−4.0, 2.5, 0.4); for the lobule VI of the cerebellum, (−6.0, 0.0, 0.2-0.25) and (−6.5, 0.0, 0.2-0.25). The window and headplate were adhered to the skull using dental acrylics. Mice were treated with pre-operative dexamethasone (20 mg/kg) and post-operative carprofen (5 mg/kg) to reduce inflammation and were allowed to recover for at least two weeks before imaging.

### Acute slice preparation

Mice with neurons transfected via stereotaxic AAV injection (0.65 mm lateral, 1mm anterior to bregma, 0.35mm deep), were perfused with ice-cold, gassed aCSF. The brain was then resected and sliced in a 10-degree tilted coronal plane using a vibratome (Leica VT1200s) in an ice-cold, gassed choline-cutting solution containing (in mM): 110 choline chloride, 25 NaHCO_3_, 25 D-glucose, 2.5 KCl, 7 MgCl_2_, 0.5 CaCl_2_, 1.25 NaH_2_PO_4_, 11.5 sodium ascorbate, and 3 sodium pyruvate. The slices were then incubated in carbogen-gassed aCSF at 35°C for 30 minutes and subsequently kept at room temperature for up to 12 hours.

### Electrophysiology

Layer 5 pyramidal neurons in acute slices of the motor cortex were recorded using the whole-cell patch-clamp technique. Transfected and adjacent (within 100 µm) untransfected neurons were recorded sequentially. Voltage- and current-clamp recordings were performed using a MultiClamp 700B amplifier (Molecular Devices) controlled by custom software written in MATLAB. Electrophysiological signals were filtered at 2 kHz before being digitized at 20 kHz. Slices were perfused at 33-35 degrees Celsius with gassed aCSF containing 2 mM CaCl_2_ and 2 mM MgCl_2_. Recording pipettes (3–5 MΩ), were pulled from borosilicate glass (G150F-3; Warner Instruments) using a model P-1000 puller (Sutter Instruments). Series resistances were 8–26 MΩ. The internal solution contained (in mM): 128 K-gluconate, 4 MgCl^2^, 10 HEPES, 10 Na-phosphocreatine, 3 Na-L-ascorbate, 0.4 Na-GTP, 4 Na-ATP, 1 EGTA and 4 QX-314 bromide with an osmolarity of 303 mOsmol/kg and pH 7.24 adjusted with CsOH. Voltages were not corrected for the liquid junction potential.

Monosynaptic excitatory postsynaptic currents (EPSCs) were evoked by extracellular stimulation of axons in the motor cortex using a theta-glass (20-50 µm tip) electrode filled with aCSF and positioned ∼200 µm ventral from somata. Short (0.1 ms), weak (25-500 µA) pulses were applied using an A365 stimulus isolator (World Precision Instruments). Direct activation of the recorded neuron was prevented by blocking action potentials with 4 mM QX-314 in the internal solution. GABA_A_ currents were blocked by adding 50–100 µM picrotoxin in the bath. In voltage-clamp mode, cells were first held at −70 mV to record AMPA currents and then at +55 mV to record NMDA currents. AMPA/NMDA ratios were calculated as the peak amplitudes of currents evoked at −70 mV divided by the average amplitudes of currents evoked at +55 mV, 40- 50 ms after stimulation. Paired-pulse ratios (PPRs) were calculated as the ratio between the amplitudes of two EPSCs (2nd peak/1st peak) evoked by electrical stimuli separated by 70 ms while holding the cell at −70 mV. 10–20 trials were recorded with an interstimulus interval of 10 s.

Miniature EPSCs were recorded in the presence of 1 μM TTX, 10 μM CPP, and 100 μM picrotoxin (to block action potentials, NMDA receptors, and GABA_A_ receptors, respectively). mEPSCs were detected as negative deflections that exceeded 5 times the median absolute value of the smoothed first derivative of the current traces. Input resistance was calculated from a test pulse in voltage-clamp mode. In current-clamp mode, instantaneous firing rates were calculated as the inverse of the first interspike interval following current injection, and steady-state firing rates were calculated as the inverse of the average interspike interval in the last half of a 1s current injection.

### Two-photon and 2pFLIM imaging

The *in vitro* two-photon microscope was built as previously described^24^ and the *in vivo* two-photon microscope was built based on the open-access design of the Modular In vivo Multiphoton Microscopy System (MIMMS) from Howard Hughes Medical Institute Janelia Research Campus (https://www.janelia.org/open-science/mimms). FLIM capacity was added as previously described^8,23^. Briefly, A photodiode (Thorlabs FDS010) was added to detect the arrival of the laser pulses. The output of a fast photomultiplier tube (Hamamatsu H7422PA-40, H10769PA-40, or H10770A-40) was compared with laser pulse timing using a TCSPC-730 (Becker and Hickl), or TimeHarp 260 (PicoQuant) time-correlated single photon counting board. Experiments using TCSPC-730 were controlled by the ScanImage software^24^ (Vidrio) integrated with a modified add-on called FLIMimage, which was written in MATLAB and was kindly provided by Dr. Ryohei Yasuda. Experiments using TimeHarp were controlled by FLIMage (Florida Lifetime Imaging). Fluorophores were excited with a pulsed 80 MHz Titanium-Sapphire laser at the following wavelengths: (850 nm for CKAR and CKAR2; 920 nm for ITRACK and IDOCKS; and 960 for CKAR3, mutCKAR3, and KCP). The fluorescence emission for these were unmixed using a dichroic mirror and band-pass filters. Specifically, Semrock FF511-Di01, Semrock FF01-483/32 and Semrock FF01-550/49 were used for CKAR, CKAR2, and additional CKAR variants and for ratiometric imaging of KCP. The Chroma 565DCXR dichroic with Semrock FF01-630/92 and Chroma HQ510/70 barrier filters was used for ITRACK, IDOCKS and CKAR3, and for lifetime imaging of KCP.

During imaging experiments, animals were head-fixed and placed on a one-dimensional treadmill that was fixed except during locomotion-related experiments. CKAR3 and mutCKAR3 were excited at 920 nm. Enforced-running experiments were carried out using a motorized treadmill approximately 5 cm/s at the indicated time. For the pharmacological experiment, the mouse was first exposed to the enforced running described above and then administered drugs via subcutaneous (s.c.) injection 30 minutes before the enforced running. JNJ16259685 (JNJ: Tocris) was dissolved in DMSO at 10 mg/mL and then diluted in saline to final concentration of 1 mg/mL. Prazosin (przn: thermos scientific) and scopolamine (scop: Tocris) were dissolved in saline at final concentration of 250 *μ*g/mL. JNJ, prazosin and scopolamine were subcutaneously administered at the dose of 4 mg/kg, 1 mg/kg and 1 mg/kg, respectively. Saline was administered as the vehicle control.

### Image analysis

Data analyses were performed using custom software written in MATLAB. In particular for 2pFLIM analyses, we used a software suite called FLIMview written in MATLAB^25^. Where appropriate, regions of interest were drawn to isolate HEK cell, somatic or dendritic signals from contamination by background photons or adjacent cells/structures. For ratiometric imaging, individual channels were background subtracted based on a nearby background ROI before calculating the ratio. Fluorescence lifetime was approximated by mean photon emission time τ:

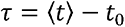

such that t reflects the emission time of individual photons in the measurement window and t_0_ reflects the timing of laser pulses. t_0_ is a fixed property of a given hardware configuration and is measured separately under ideal imaging conditions. Because the measurement window (t_w_) is finite (≤ 10 ns passing t_0_ due to laser pulse repetition and single photon counting hardware properties), the measured τ (τ_apparent_) is slightly smaller than the real value. For example, for a τ of 2.0 ns and a t_w_ of 9.0 ns, the τ_apparent_ is 1.90 following the equation below:

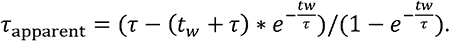

### Analysis of *in vivo* data

The basal lifetime of each cell was calculated as the mode of the distribution estimated by Kernel Density Estimation on the 5-minute baseline. For Fig. 3e and i, Pearson’s correlation coefficient was used to represent the correlation of basal lifetimes and area under curve (AUC) during enforced running within the same cells in the M1 cortex imaged 5–8 days apart. For Fig. 3j, hierarchical clustering using Ward’s method was performed on the data normalized to the peak response of each responder cell in Fig. 3h to enforced running. The optimal number of clusters was determined based on the highest mean silhouette score (ED Fig. 7a). For Fig. 3k and l, the data were smoothed using a moving average filter with a window size of 5, and peaks and their timing were detected as 3 times more than the standard deviation of the baseline in running sessions. For Fig 3m, the on- and off-kinetics of each cluster were calculated using one-phase decay with least squares regression from data normalized to the first and last peaks of each cell, respectively. For Fig 3o, the response to the enforced running after drug administration was normalized to the control response prior to the drug administration.

### Data presentation and statistical analysis

Quantification and statistical tests were performed using custom software written in MATLAB and GraphPad Prism 10. Averaged data are presented as mean ± s.e.m., unless noted otherwise. All measurements were taken from different cells. “n” refers to the number of cells unless noted. In HEK cell experiments, most (∼70%) experiments measured two cells simultaneously per coverslip. In a few specific experiments, up to ten cells per coverslip were imaged. In cultured and acute slice experiments, all neurons except a pair came from different slices. All experiments were repeated for at least two, typically more, independent transfections (*in vitro*) or mice (*in vivo*). Unless otherwise noted, the Shapiro Wilk test was used to determine if data were normally distributed, and the Bartlett test was used to determine if variances were equal. When the assumptions of normality and homoscedasticity were met, a two-tailed Student’s t-test or one-way ANOVA followed by a Bonferonni post-hoc test were used. When the assumption of normality was not met, Wilcoxon signed-rank test, Mann-Whitney U test and Kruskal-Wallis test followed by Dunn’s post hoc test were used for paired and unpaired comparison. In all figures, *: p < 0.05, **: p < 0.01, and ***: p < 0.001.

## Acknowledgements

We thank Priscilla Ambrosi for preliminary efforts in electrophysiology that are not included in the manuscript. This work was supported by two NIH BRAIN Initiative awards R01NS104944 and RF1MH130784 (H.Z. and T.M.), two NINDS R01 grants R01NS081071 (T.M.) and R01NS127013 (H.Z.), and a NIDA F30 grant (F30DA057838).

## Author Contributions

HZ conceived the project and generated the initial variants. TY, LBE, TM, and HZ designed the experiments. LBE and HZ performed *in vitro* imaging experiments. TY performed *in vivo* experiments with initial guidance from LM and assistance from HZ and LBE. JAJ performed electrophysiological recordings. MZ prepared and maintained cultured slices and mouse husbandry. TY, LBE, JAJ, and HZ analyzed the data. TY, LBE, JAJ, TM, and HZ wrote the manuscript, with edits and comments by all authors. TM and HZ secured the funding and supervised the project.

## Data availability

All previously unpublished sensor constructs and their corresponding sequences will be deposited to Addgene (http://addgene.org). Source data will be provided with this paper.

## Code availability

Custom MATLAB codes will be made available upon request.

## Extended Data Figures and Table

**Extended Data Figure 1.**
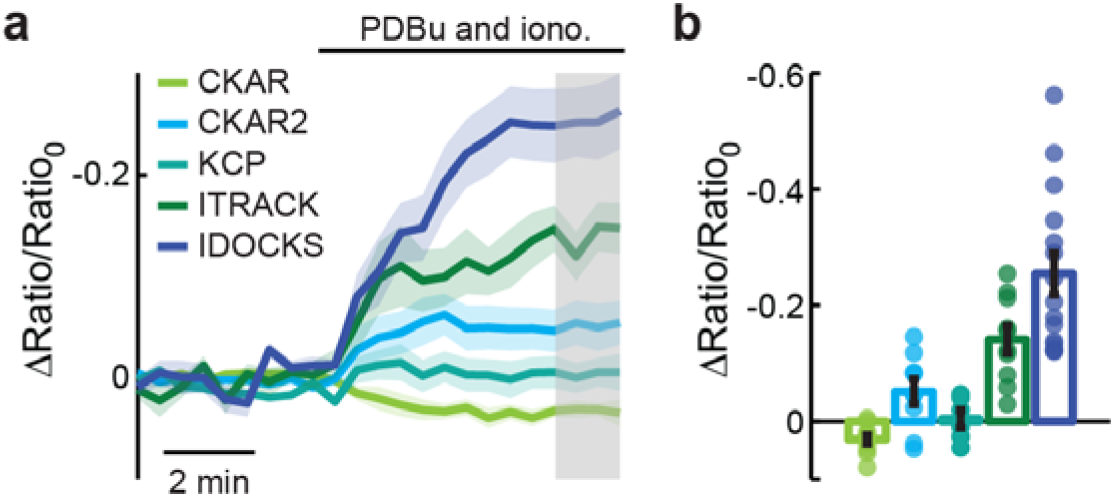
Comparison of the ratiometric responses of existing PKA sensors. **a** & **b**, Averaged ratiometric response traces (**a**) and quantification in the grey time interval (**b**) of existing PKC sensors responding to PDBu and ionomycin. Responses were measured as the normalized ratio of the donor to acceptor FP intensities. From left to right, n (cells/coverslip) = 11/4, 9/4, 6/3, 10/5, 15/6. Dark lines and shaded areas indicate mean and s.e.m, respectively. Bars indicate mean and error bars indicate s.e.m.

**Extended Data Figure 2.**
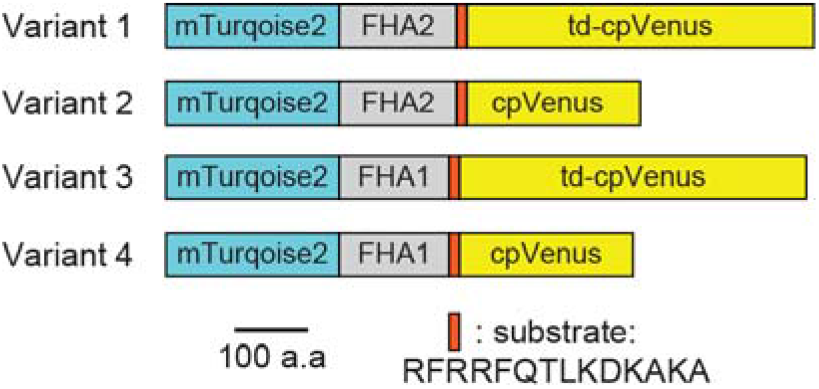
Schematic of tested CKAR3 variants. FHA1 and FHA2: Forkhead-associated domains 1 and 2, respectively.

**Extended Data Figure 3.**
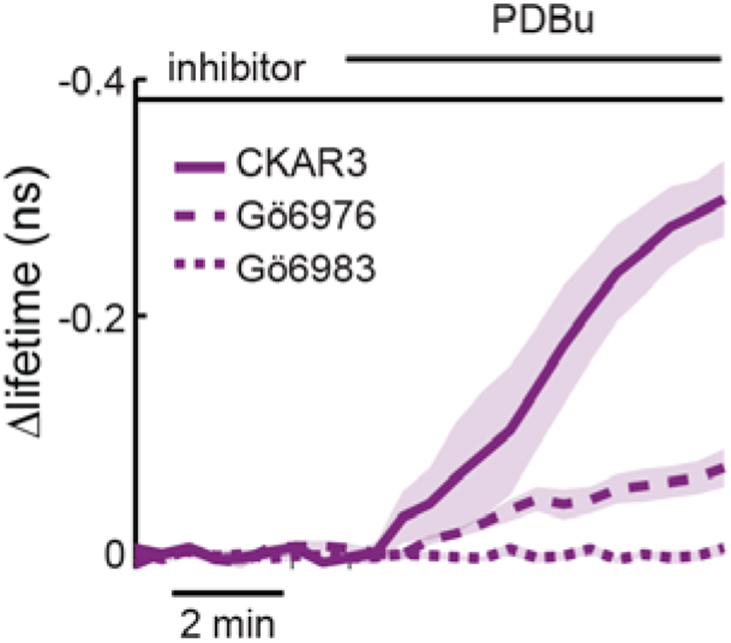
PKC inhibitors blocked CKAR3 response to PDBu. Average response of CKAR3 to PDBu after pre-treatment with 1 μM Gö6976 or Gö6983. n (cells/coverslips) = 14/4, 7/3, 8/3 for CKAR3 with no inhibitor, with Gö6976 or Gö6983, respectively. Dark line and shaded area indicate mean and s.e.m, respectively. Bars indicate mean and error bars indicate s.e.m.

**Extended Data Figure 4.**
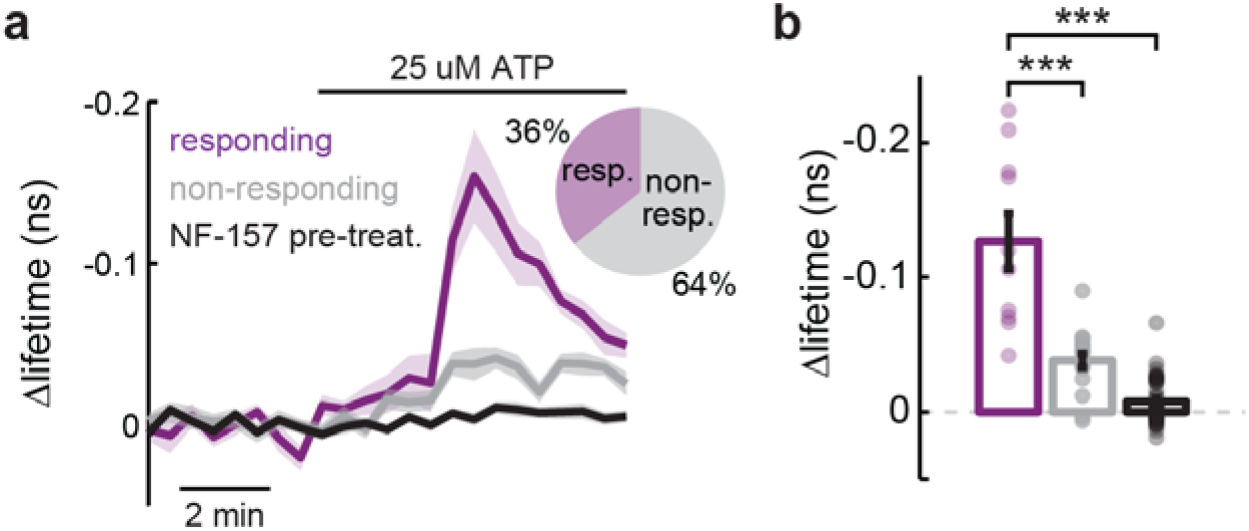
ATP application causes PKC activation through P_2_Y_11_ in a subset of cultured HEK cells. **a**, Average response of CKAR3 to 25 μM ATP application with and without pre-treatment with 1 μM NF-157, a P_2_Y_11_ antagonist. Responding cells (threshold 2x baseline s.d.) are shown in purple, non-responding in grey, and pretreated cells in black with quantification in (**b**). From left to right, n (cells/coverslips) = 10/3, 18/3 and 43/5. ***: p< 0.001 (0 for both comparisons), dF = 2, one-way ANOVA (p = 9x10^-17^), Bonferroni post-hoc test. Dark line and shaded area indicate mean and s.e.m, respectively. Bars indicate mean and error bars indicate s.e.m.

**Extended Data Figure 5.**
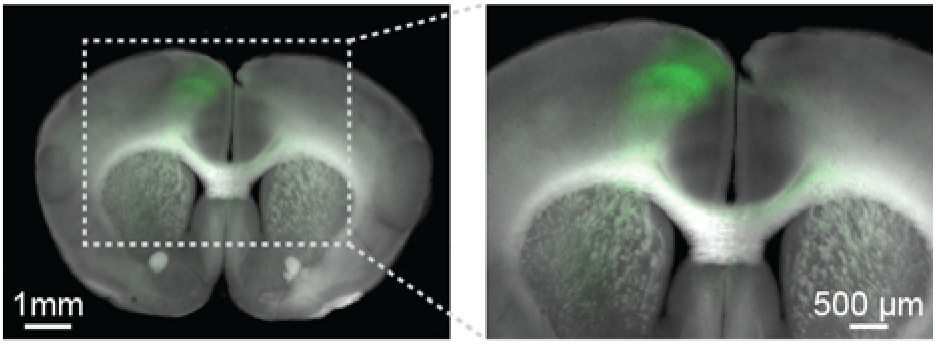
Representative image of CKAR3 expression in the M1 cortex. CKAR3 was expressed via AAV injection under a synapsin promoter.

**Extended Data Figure 6.**
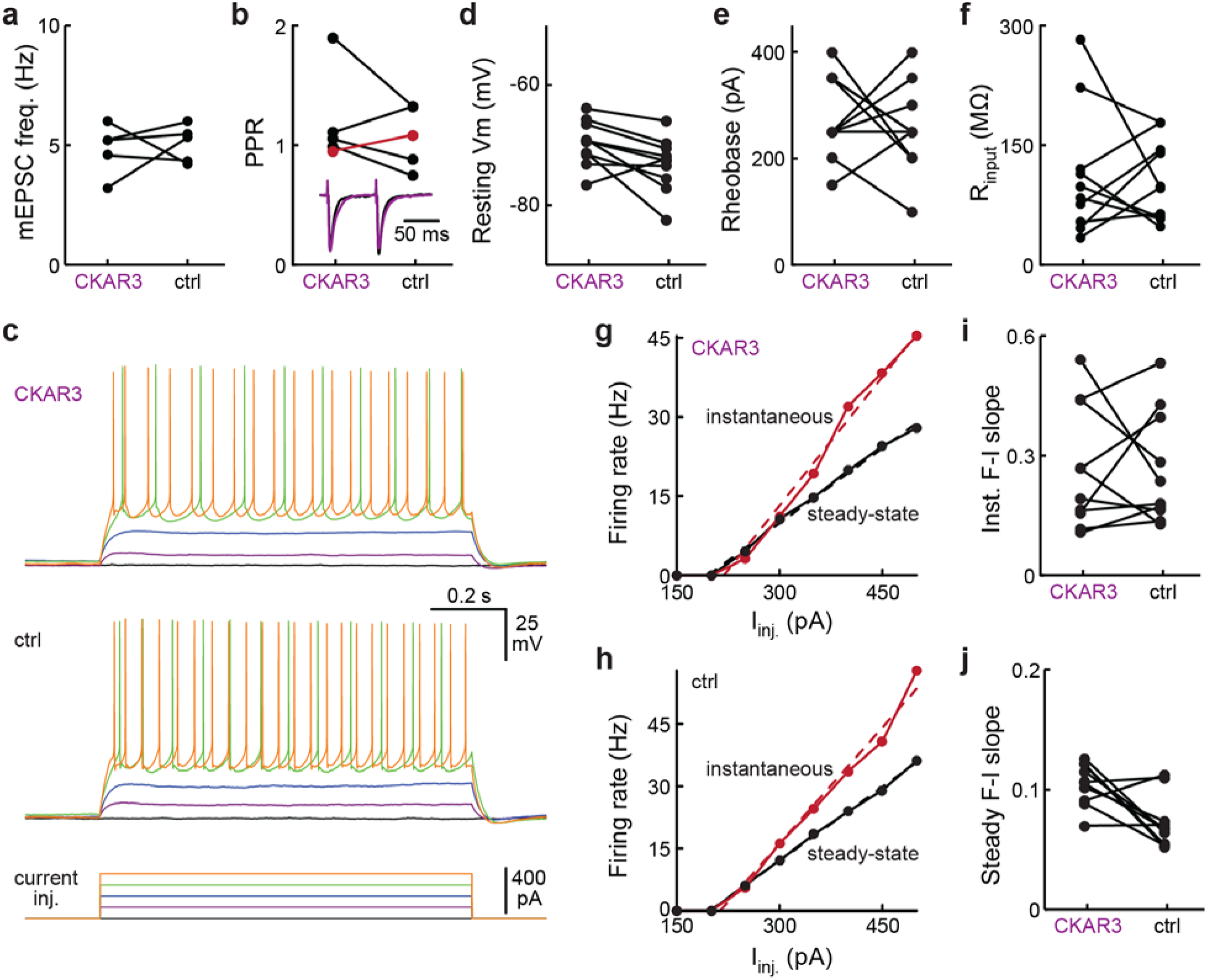
Additional electrophysiological characterizations of CKAR3 expressing neurons. **a**, mEPSC frequency of transfected L5 pyramidal neurons compared to adjacent untransfected controls. n (pairs/mice) = 5/4, p = 0.74, dF = 4, paired t-test. **b**, Paired-pulsed ratio. n = (pairs/mice) = 5/3, p = 0.42, signed-rank test. **c**, Membrane voltage responses of (top) an L5 motor cortical pyramidal neuron transfected with CKAR3 and (middle) an adjacent untransfected pyramidal neuron to (bottom) 1s current steps. **d**–**f**, Resting membrane potentials (**d**), rheobases (**e**), and input resistances (**f**) of transfected pyramidal neurons compared to adjacent untransfected pyramidal neurons. From left to right, n (pairs/mice) = 10/2, 10/2, 10/2; p = 0.03, 0.53, 0.80; dF = 9, 9, 9; paired t-tests. **g** & **h**, (Red) Instantaneous and (black) steady-state F-I curves of a transfected (**g**) and an adjacent untransfected (**h**) pyramidal neurons. **i** & **j**, Slopes of instantaneous (**i**) and steady-state (**j**) F-I curves of transfected compared to adjacent untransfected neurons. From top to bottom, n (pairs/mice) =10/2, 10/2; p = 0.92, 0.02; dF = 9, 9; signed-rank tests. Upon correcting for multiple comparisons, there were no significant effects of CKAR3 expression.

**Extended Data Figure 7.**
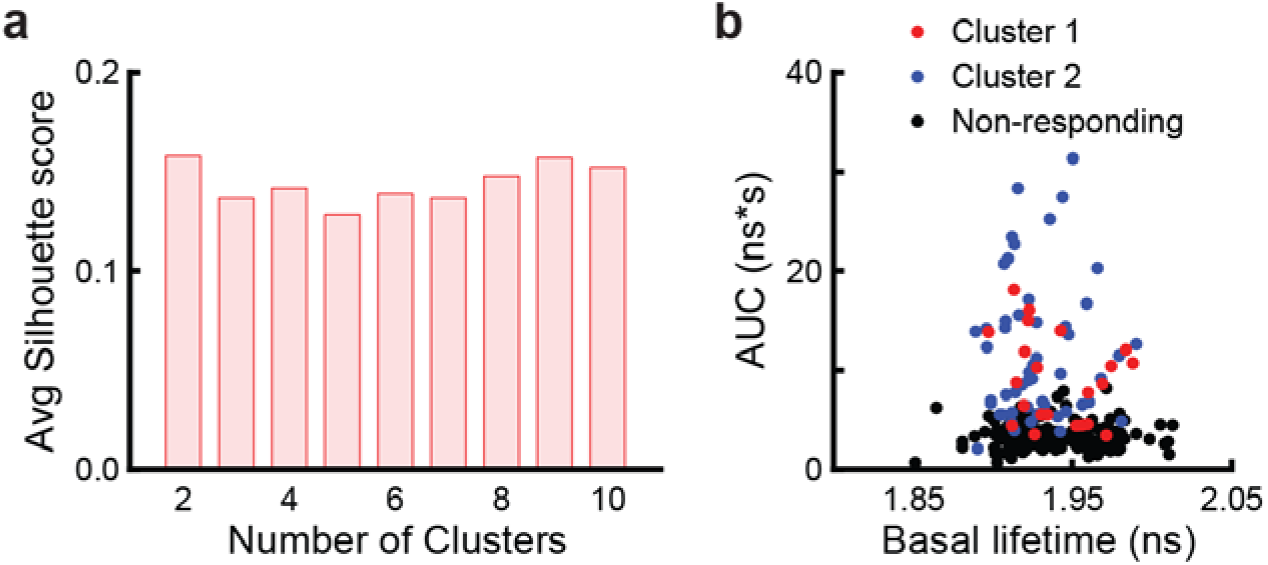
Supporting data for clustering analysis of locomotion-elicited CKAR3 responses. **a**, Averaged silhouette score for each potential number of clusters. **b**, The relationship between basal lifetime and response magnitude for each cluster. *n* (cells) = 22 for cluster 1, *n* = 52 for cluster 2 and 166 for non-responders from 5 mice.

